# Inducible degradation of dosage compensation protein DPY-27 facilitates isolation of *Caenorhabditis elegans* males for molecular and biochemical analyses

**DOI:** 10.1101/2022.01.27.478040

**Authors:** Qianyan Li, Arshdeep Kaur, Benjamin Mallory, Sara Hariri, JoAnne Engebrecht

## Abstract

Biological sex affects numerous aspects of biology, yet how sex influences different biological processes has not been extensively studied at the molecular level. *Caenorhabditis elegans*, with both hermaphrodites (functionally females as adults) and males, is an excellent system to uncover how sex influences physiology. Here, we describe a method to isolate large quantities of *C. elegans* males by conditionally degrading DPY-27, a component of the dosage compensation complex essential for hermaphrodite, but not male, development. We show that germ cells from males isolated following DPY-27 degradation undergo meiosis and spermiogenesis like wild type and are competent to mate and produce viable offspring. We demonstrate the efficacy of this system by analyzing gene expression and performing affinity pull-downs from male worm extracts.

## Introduction

In metazoans, sex has evolved multiple times and influences most biological processes. Gonochoristic species have two biological sexes, defined by the production of differentiated gametes: sperm (males) and ova (females). Somatic tissues also display sexually dimorphic features, the most obvious being those important for mating. Further, studies in mammals have highlighted the impact of sex on physiological processes including metabolism, cardiac and neuronal functions (Miller 2014). The molecular mechanisms underlying how biological processes are modulated by sex remain largely unknown.

In addition to gonochorism, other reproductive strategies exist. For example, hermaphrodism is common in many species including snails, worms, echinoderms, fish and plants, where both sperm and ova are produced in the same organism. The nematode *Caenorhabditis elegans* is an androdioecious species with both hermaphrodites and males and has been an excellent model to study sex-specific morphological adaptations and production of sperm and ova. In *C. elegans* hermaphrodites (*XX*), the first wave of germ cells undergoes spermatogenesis; as adults, hermaphrodites exclusively produce ova and thus are functionally female (Hubbard and Greenstein 2000). Males (*XO*) arise spontaneously at a low frequency (∼0.2%) because of meiotic chromosome nondisjunction and exclusively produce sperm. The morphological features that differentiate the *C. elegans* male from the hermaphrodite arise during postembryonic development. Most prominent is the tail structure and associated male-specific neuronal wiring required for mating. Males can be propagated by crossing with hermaphrodites, which will preferentially use male sperm to fertilize ova, leading to a 1:1 hermaphrodite:male ratio in the offspring (LaMunyon and Ward 1995, 1998; Ward and Carrel 1979).

*C. elegans* is a facile genetic model due in part to its facultative hermaphroditic lifestyle, which greatly simplifies genetic analysis. The wealth of genetic mutants available and in-depth understanding of how sex is determined, facilitates mechanistic studies of how sex affects biological processes. For example, *fog-2* loss-of-function mutants block spermatogenesis specifically in hermaphrodites, leading to true female worms (Schedl and Kimble 1988). These mutant worms have been used to distinguish sperm versus ova contributions to euploid progeny (Jaramillo-Lambert et al. 2010; Li, Hariri, and Engebrecht 2020; Checchi et al. 2014). Several genes required for spermatogenesis (*spe* genes) have been identified. Some *spe* genes are required for spermatogenesis in both hermaphrodites and males (Nishimura and L’Hernault 2010). Conditional depletion of one of these, *spe-44*, has been developed for mating and longevity studies (Kasimatis, Moerdyk-Schauwecker, and Phillips 2018). Interestingly, the *spe-8* group is specifically required for activation of hermaphrodite, but not male, sperm (L’Hernault, Shakes, and Ward 1988). Additionally, mutations in genes that are important for *X* chromosome disjunction in meiosis leads to the Him (<u>H</u>igh incidence of <u>m</u>ales) phenotype, such that up to 40% of self-progeny are males (Hodgkin, Horvitz, and Brenner 1979).

*C. elegans* has prominent gonads where germ cells are organized in a linear assembly line fashion and reproduces prolifically, making *C. elegans* an outstanding system to investigate meiosis and fertilization. To date, most meiotic studies have focused on oogenesis in hermaphrodites due to the ease of isolating meiotic mutants and performing molecular analyses. A few studies focusing on male meiosis have revealed that while the basic processes are similar, the regulation of meiosis is distinct in spermatogenesis versus oogenesis (Checchi et al. 2014; Jaramillo-Lambert et al. 2010; Kurhanewicz et al. 2020; Jaramillo-Lambert et al. 2007; Shakes et al. 2009; Li, Hariri and Engebrecht, 2020). However, a complete understanding of how sex influences meiosis, gametogenesis, and biology more generally, is still lacking.

While hermaphrodites are easy to propagate and collect in large numbers, it has been more difficult to propagate and collect large numbers of males. Most strategies rely on maintaining mated cultures (∼50% males) or using *him-8* mutants (∼ 40% males) (Hodgkin, Horvitz, and Brenner 1979) and then manually picking males. Alternatively, males can be separated from hermaphrodites using filters, which takes advantage of the different size of adult hermaphrodites (1mm × 80μm) and males (0.8mm × 50 μm) (Chu et al. 2006). However, this requires synchronized cultures as larvae are smaller and will filter with males regardless of sex. Additionally, filtering requires extensive labor and in our hands is not very efficient, making it difficult to collect large quantities required for biochemical analyses.

Here we describe a new method to isolate relatively pure populations of males in large numbers. This method takes advantage of the inducible degradation of DPY-27, a component of the dosage compensation complex (DCC) (Plenefisch, DeLong, and Meyer 1989). In *C. elegans*, the DCC down regulates gene expression from the two *X* chromosomes in hermaphrodites such that the overall level is comparable to the expression from the single *X* chromosome in males. Consequently, the DCC is essential for embryonic development in hermaphrodites but not in males. Worms defective for the DCC are therefore hermaphrodite-specific lethal (Meyer 2005). Using the auxin-inducible degradation system that has been adapted for *C. elegans* (Ashley et al. 2021; Zhang et al. 2015), we fused a degron tag to DPY-27 and constructed strains also expressing TIR1 in the *him-8* mutant background, which produces male self-progeny (Hodgkin, Horvitz, and Brenner 1979). We show that males collected after auxin treatment are proficient for meiosis, spermatogenesis, and mating. We demonstrate the effectiveness of this method by analyzing gene expression and performing affinity pull-downs followed by mass spectrometry from male worm extracts.

## Materials and Methods

### Genetics

*C. elegans* var. Bristol (N2) was used as the wild-type strain. The following strains were constructed:

JEL1197 *sun-1p::TIR1 II; dpy-27::AID::MYC (xoe41) III; him-8(me4) IV*

JEL991 *sun-1p::TIR1 II; dpy-27::AID::MYC(xoe41) brd-1::gfp::3xflag(xoe14) III; him-8(me4) IV*

JEL1217. *sun-1p::TIR1 II; gfp(glo)::3xflag::cosa-1(xoe44) dpy-27::AID::MYC (xoe41) III; him-8(me4) IV*

JEL1214. *mex-5p::TIR1 I; dpy-27::AID::MYC (xoe41) III; him-8(me4) IV*

*sun-1p::TIR1* and *mex-5p::TIR1* strains were graciously provided by Jordan Ward (UCSC) (Ashley et al. 2021). Some nematode strains were provided by the Caenorhabditis Genetics Center, which is funded by the National Institutes of Health National Center for Research Resources (NIH NCRR). Strains were maintained at 20°C.

### CRISPR-mediated generation of alleles

*dpy-27::AID::MYC(xoe41)* and *gfp(glo)::3xflag::cosa-1(xoe44)* were generated using the co-CRISPR method as described (Paix et al. 2015). <u>G</u>erm<u>L</u>ine <u>O</u>ptimized (GLO) GFP sequence was used to enhance germline expression and prevent potential silencing (Fielmich et al. 2018). Guide sequence, repair template and genotyping primers are provided in Supplemental File 1. Correct editing was verified by Sanger sequencing. Worms generated by CRISPR were outcrossed a minimum of four times.

### Auxin treatment to generate male cultures

Synchronized L1 larvae were placed on NGM plates containing 1-2mM auxin (Naphthaleneacetic Acid; K-NAA; PhytoTech #N610) and maintained at 20°C until adulthood. Worms were washed off plates and bleached with 5N NaOH and 5% bleach (Porta-de-la-Riva et al. 2012). Embryos were collected by centrifugation at 1300g for 1min and washed 3x with M9 buffer supplemented with auxin. Embryos were transferred to a new tube with M9 buffer + auxin and kept on a rocker for 24hr. L1 larvae together with unhatched embryos were washed with M9 buffer and plated onto NGM plates without auxin. Approximately two and half days later, viable worms were quantified for % male and used for downstream analyses. A detailed protocol for large-scale collection of male worms is provided in Supplemental File 2.

### Cytological analyses

Immunostaining of germ lines was performed as described (Jaramillo-Lambert et al. 2007) except slides were fixed in 100% ethanol instead of 100% methanol for direct GFP fluorescence of GFP::COSA-1. Rabbit anti-RAD-51 (1:10,000; cat #2948.00.02; SDIX; RRID: AB_2616441) and secondary antibodies Alexa Fluor 594 donkey anti-rabbit IgG from Life Technologies were used at 1:500 dilutions. DAPI (2μg/ml; Sigma-Aldrich) was used to counterstain DNA.

Collection of fixed images was performed using an API Delta Vision Ultra deconvolution microscope equipped with an 60x, NA 1.49 objective lens, and appropriate filters for epi-fluorescence. Z stacks (0.2 μm) were collected from the entire gonad. A minimum of three germ lines was examined for each condition. Images were deconvolved using Applied Precision SoftWoRx batch deconvolution software and subsequently processed and analyzed using Fiji (ImageJ) (Wayne Rasband, NIH).

RAD-51 foci were quantified in germ lines of age-matched males (18-24hr post-L4). Germ lines were separated into the transition zone (leptotene/zygotene), as counted from the first and last row with two or more crescent-shaped nuclei, and pachytene, which was further divided into 3 equal parts: early, mid and late pachytene. RAD-51 foci were quantified from half projections of the germ lines. The number of foci per nucleus was scored for each region.

GFP::COSA-1 foci were quantified from deconvolved 3D data stacks; mid-late pachytene nuclei were scored individually through z-stacks to ensure that all foci within each individual nucleus were counted.

Spermiogenesis was monitored as described by releasing sperm into sperm medium (50mM HEPES, 25mM KCl, 45mM NaCl,1mM MgSO4, 5mM CaCl2, 10mM Dextrose; pH 7.8) in the absence and presence of 20μg Pronase E (MedChemExpress HY-114158A) and imaged under DIC on a Zeiss Axioscope (Singaravelu et al. 2011).

### Viability of male-sired progeny

A single *fog-2(q71)* female was mated with 3 males of indicated genotypes on small *E. coli* OP50 spots. The mated female was transferred to new plates every 24hr. Progeny viability was determined over 3 days by counting eggs and hatched larvae 24hr after removing the female and calculating percent as larvae/(eggs + larvae). The progeny of a minimum of 8 mated females were scored.

### RT-PCR

Total RNA was isolated from 50-100μl of packed worms from indicated genotypes/conditions using the RNeasy Mini Kit (Qiagen, Catalog #74104) and QIAshredder (Qiagen, Catalog #79654). 1μg of RNA was converted to cDNA using SuperScript III First-Strand Synthesis System for RT-PCR (Invitrogen, Catalog #18080-051) primed with Oligo(dT)_20_. qPCR reactions were prepared with SsoAdvanced Universal SYBR Green Supermix (Bio-Rad, Catalog #1725271) using cDNA and the following primers (final concentration, 400 nM): *ama-1* (Housekeeping) Fwd: 5’-GACGAGTCCAACGTACTCTCCAAC-3’, Rev: 5’-TACTTGGGGCTCGATGGGC-3’; *vit-2* (female-enriched) Fwd: 5’-GCCAGAAGAACCAGAGAAGCC-3’, Rev: 5’-TGTTGTTGCTGCTCGACCTC-3’; *sncp-1*.*3* (male-enriched) Fwd: 5’-TCCTTCATGCGAATGACCCG-3’, Rev: 5’-GCGCTTTGAATCTACCCAGC-3’; Cq values were determined for each primer pair and normalized to the *ama-1* control. The fold change between – or + auxin and wild type was analyzed using the 2^-ΔΔCt^ method. Raw Cq values and calculations are provided in Supplemental File 3.

### Pull-down assays

Male worms were collected following auxin treatment as described above and resuspended in H100 lysis buffer (50mM HEPES, pH 7.4, 1mM EGTA, 1mM MgCl2, 100mM KCl, 10% glycerol, 0.05% NP-40) + protease inhibitors (Complete Ultra Tablets, Mini Protease Inhibitor Cocktail; Roche #05892791001 and 1mM PMSF; Sigma). Worms were allowed to settle at the bottom of the tube and the lysis buffer was removed until there was 0.5-1ml buffer covering the worm pellet. Worms were resuspended and flash frozen as “worm popcorns” by pipetting into liquid nitrogen. Worm popcorns were ground into fine powder using a SPEX SamplePrep 6970 FreezerMill and immediately stored at -80°C.

Approximately 1ml of worm powder was thawed on ice and brought to 4ml with H100 + protease inhibitors. The lysate was passed through a glass tissue grinder multiple times on ice and then centrifuged at 13000 rpm for 20min at 4°C to remove insoluble debris. 50μl of Anti-FLAG M2 magnetic beads (Millipore Sigma M8823) were prepared for each pull-down assay by washing with 1ml H100 + protease inhibitors 4x using a magnetic rack separator. The soluble fraction was incubated with the washed beads for 3hr with constant rotation at 4°C. Beads were washed with 1ml lysis buffer + protease inhibitors 4x and then washed in lysis buffer without protease inhibitors 7x. 1/10 of beads were analyzed by immunoblot to check efficiency of pull-down and the remaining processed for mass spectrometry.

### Mass spectrometry analyses

Processing and proteomic profiling was performed at the University of California Davis Proteomics Core Facility (https://proteomics.ucdavis.edu). Protein samples on magnetic beads were washed 4x with 200μl 50mM Triethyl ammonium bicarbonate (TEAB) for 20min each at 4°C with shaking. 2μg of trypsin was added to the bead/TEAB mixture and the samples were digested ON at 4°C with shaking at 800 rpm. The supernatant was then removed, and the beads washed once with enough 50mM ammonium bicarbonate to cover. After 20min with gentle shaking the wash was removed and combined with the initial supernatant. The peptide extracts were reduced in volume by vacuum centrifugation and a small portion of the extract was used for fluorometric peptide quantification (Thermo scientific Pierce). One microgram was loaded for each LC-MS analysis.

LC MS/MS was performed on an ultra-high-pressure nano-flow Nanoelute (Bruker Daltonics) at 40°C with a constant flow of 400nl/min on a PepSep 150umx25cm C18 column (PepSep, Denmark) with 1.5 μm particle size (100 Å pores) and a ZDV captive spray emitter (Bruker Daltonics). Mobile phases A and B were water with 0.1% formic acid (v/v) and 80/20/0.1% ACN/water/formic acid (v/v/vol), respectively. Peptides were separated using a 30min gradient. Eluting peptides were then further separated using TIMS (trapped ion mobility spectrometry) on a Bruker timsTOF Pro 2 mass spectrometer. Mass spectrometry data was acquired using the dda PASEF method (https://pubmed.ncbi.nlm.nih.gov/30385480/). The acquisition scheme used was 100ms accumulation, 100ms PASEF ramp (at 100% duty cycle) with up to 10 PASEF MS/MS scans per topN acquisition cycle. The capillary voltage was set at 1700V, Capillary gas temp 200°C. The target value was set at 20,000 a.u. with the intensity threshold set at 500 a.u. The m/z range surveyed was between 100 to 1700. Precursor ions for PASEF-MS/MS were selected in real time from a TIMS-MS survey scan using a non-linear PASEF scheduling algorithm. The polygon filter (200 to 1700 m/z) was designed to cover ions within a specific m/z and ion mobility plane to select multiply charged peptide features rather than singly charged background ions. The quadrupole isolation width was set to 2 Th for m/z < 700 and 3 Th for m/z 800.

Mass spectrometry raw files were searched using Fragpipe 16.0 (Kong et al. 2017) against the UniProt *Caenorhabditis elegans*; UP000001940 database. Decoy sequences were generated, appended and laboratory contaminates added within Fragpipe. Default search settings were used. Carbamidomethylation of cysteine residues was set as a fixed modification, and methionine oxidation and acetylation of protein N termini as variable modifications. Decoy False Discovery Rates were controlled at 1% maximum using both the Peptide and Protein prophet algorithms. Label-free protein quantification was performed with the IonQuant algorithm (default settings) (Yu et al. 2020).

Search results were loaded into Scaffold (version Scaffold_5.0.1, Proteome Software Inc., Portland, OR) for visualization purposes. Proteins that contained similar peptides and could not be differentiated based on MS/MS analysis alone were grouped to satisfy the principles of parsimony. Proteins sharing significant peptide evidence were grouped into clusters. The complete list of proteins identified from the control and the unique proteins identified from the BRD-1::GFP::3xFLAG pull downs are provided in Supplemental File 4.

### Statistical analyses

Statistical analyses and figures were prepared using GraphPad Prism version 9.0 (GraphPad Software). Statistical comparisons of % males (Figure 1C), RAD-51 (Figure 2A) and GFP::COSA-1 foci numbers (Figures 2B), progeny viability (Figure 2D), and fold change in *vit-2* and *sncp-1*.*3* expression (Figure 3A) were analyzed by Mann-Whitney. Detailed descriptions of statistical analyses are indicated in figure legends.

**Figure 1.**
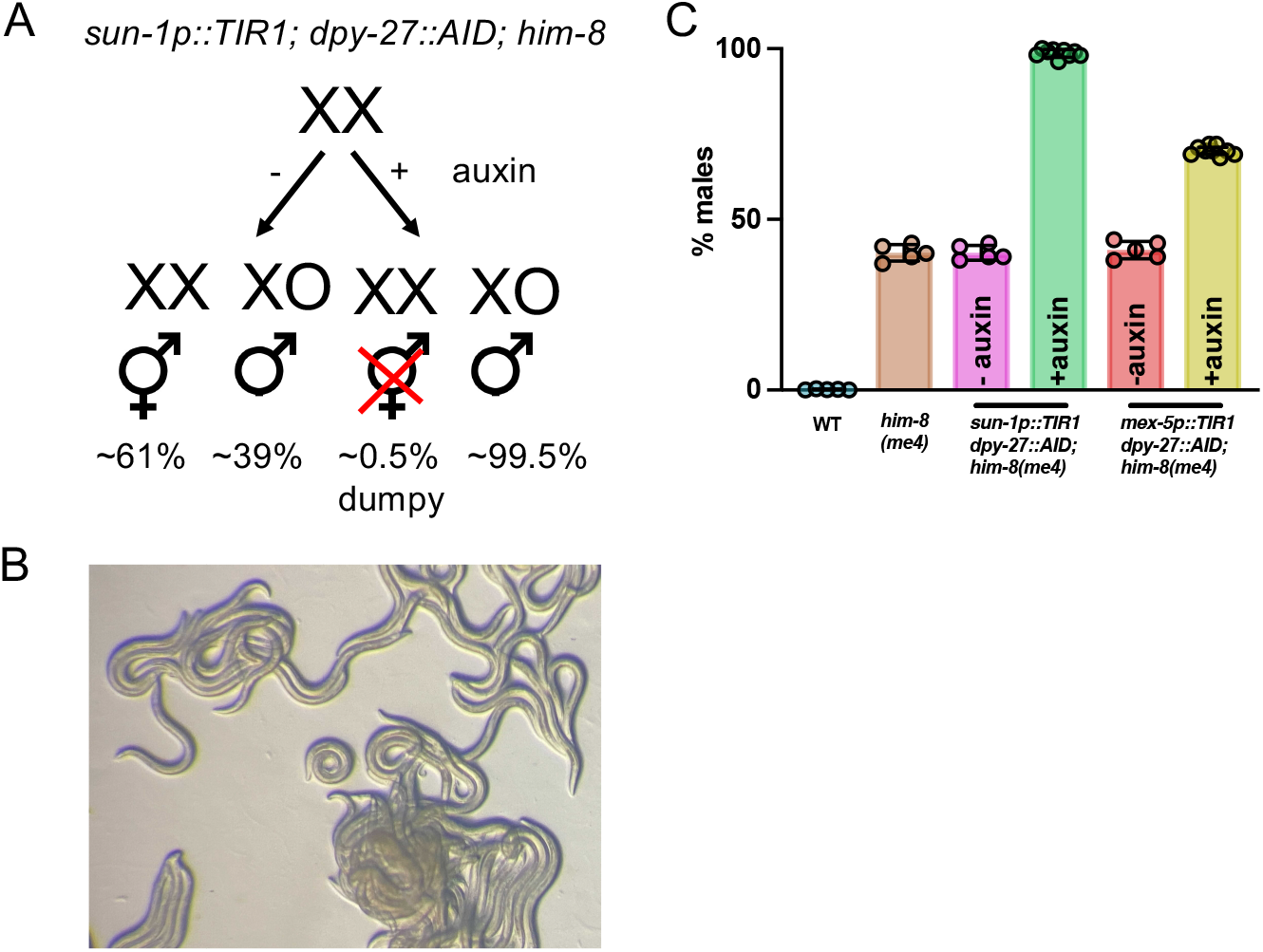
Enrichment of male worms by conditional degradation of DPY-27. A) Strategy for isolation of male cultures. B) Plate phenotype following auxin depletion of DPY-27, where greater than 98% of the worms are males. C) Quantification of male enrichment in WT, *him-8(me4), sun-1p::TIR1; dpy-27::AID; him-8(me4)* and *mex-5p::TIR1; dpy-27::AID; him-8(me4)* in the absence (-auxin) and presence (+auxin) of 1mM auxin. A minimum of 5 plates were counted for males; mean and 95% confidence intervals are shown. Statistical comparisons between + and – auxin for *sun-1p::TIR1; dpy-27::AID; him-8(me4)*: p=0.001 and *mex-5p::TIR1; dpy-27::AID; him-8(me4)*: p=0.0005 by Mann-Whitney.

**Figure 2.**
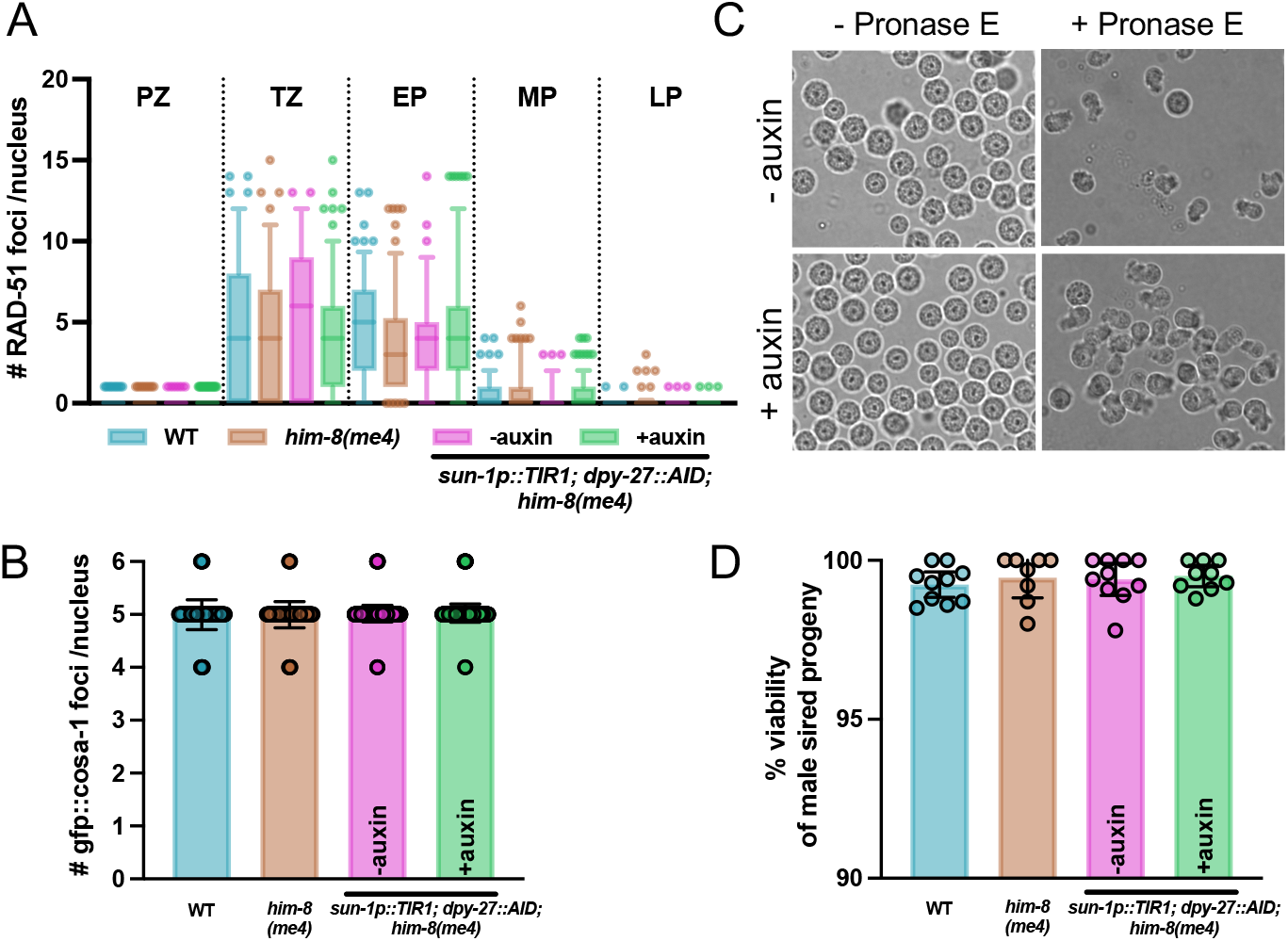
DPY-27 depletion does not affect meiosis or spermiogenesis. A) Quantification of RAD-51 in indicated regions of the germ line. Box whisker plots show number of RAD-51 foci per nucleus. Horizontal line of each box represents the median, top and bottom of each box represents medians of upper and lower quartiles, lines extending above and below boxes indicate SD, and individual data points are outliers from 5 to 95%. Comparisons by Mann–Whitney revealed no statistical differences between the strains. PZ, proliferative zone; TZ, transition zone; EP, early pachytene; MP, mid-pachytene; LP, late pachytene. Three germ lines were scored for each strain/condition. Number of nuclei scored in each region: WT, PZ = 559; TZ =136; EP = 132; MP = 175; LP = 132; *him-8(me4)*, PZ = 452; TZ = 115; EP = 134; MP = 141; LP = 135; *sun-1p::TIR1; dpy-27::AID; him-8(me4) -* auxin, PZ = 391; TZ = 126; EP = 117; MP = 155; LP = 168; *sun-1p::TIR1; dpy-27::AID*,; *him-8(me4)* + auxin, PZ = 671; TZ = 187; EP = 197; MP = 209; LP = 206. B) Number of COSA-1 foci in mid-late-pachytene in indicated strains/conditions; mean and 95% confidence intervals are shown. Comparisons by Mann-Whitney revealed no statistical differences between the strains. Number of nuclei analyzed: wild type = 189 (from 3 worms); *him-8(me4)* = 182 (from 3 worms); *sun-1p::TIR1; dpy-27::AID; him-8(me4) -* auxin = 549 (from 8 worms); *sun-1p::TIR1; dpy-27::AID; him-8(me4)* +auxin = 584 (from 8 worms). C) Micrographs of sperm isolated from *sun-1p::TIR1; dpy-27::AID; him-8(me4)* -auxin and +auxin and in the presence of Pronase E, which leads to sperm activation. D) Embryonic lethality of *fog-2(q71)* progeny sired by indicated males. Mean and 95% confidence intervals are shown. Comparisons by Mann-Whitney revealed no statistical differences between the strains/conditions. Number of crosses examined: wild type = 10; *him-8(me4)* = 8; *sun-1p::TIR1; dpy-27::AID; him-8(me4)* -auxin = 10; *sun-1p::TIR1; dpy-27::AID; him-8(me4)* +auxin = 9.

**Figure 3.**
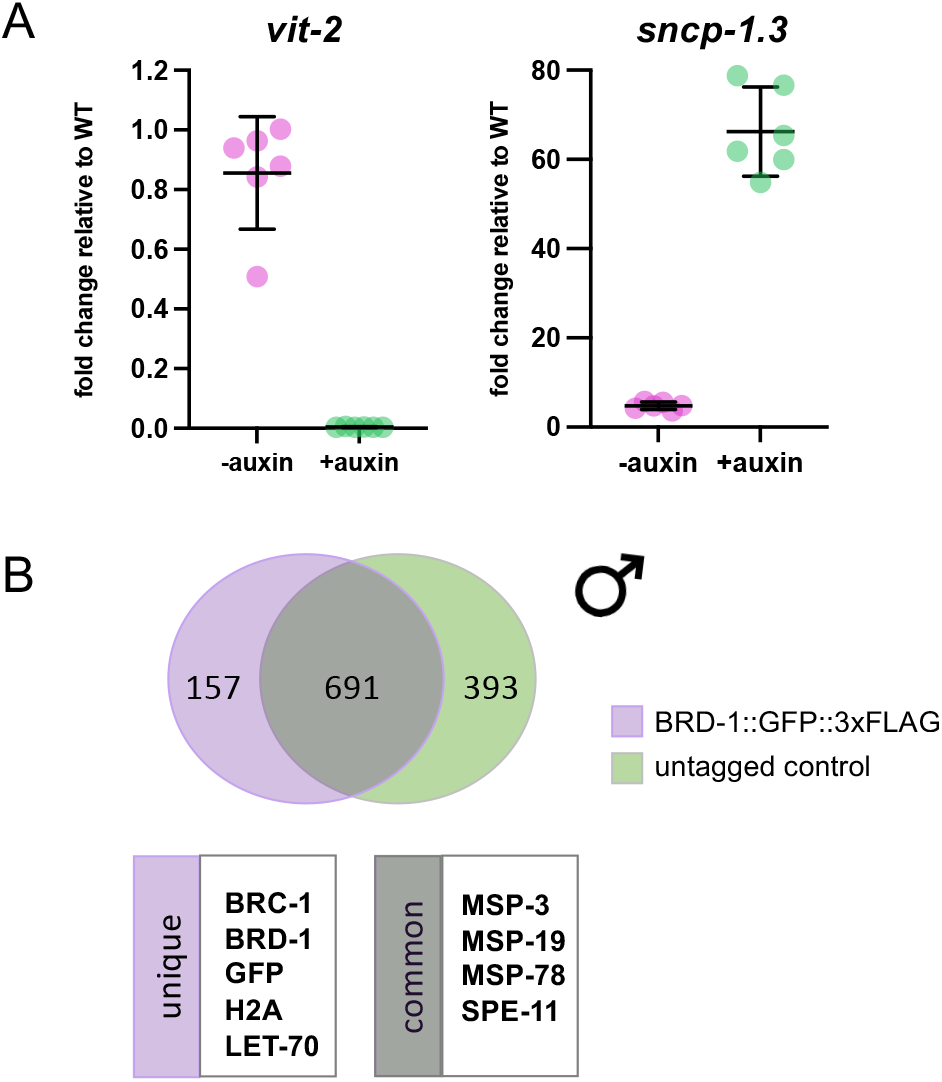
Molecular and biochemical analyses of males isolated following DPY-27 depletion. A) Fold change of mRNA expression for *vit-2* (hermaphrodite enriched) and *sncp-1*.*3* (male enriched) *in sun-1p::TIR1;dpy-27::AID; him-8(me4)* in the absence and presence of auxin relative to wild type. Fold change was determined using the 2^-ΔΔCq^ method. *ama-1* (housekeeping) mRNA levels were used as a reference. Statistical comparisons by Mann-Whitney between – and + auxin for both *vit-2* and *sncp-1*.*3*, p=0.0022. Raw Cq values and calculations are provided in Supplemental File 3. B) Mass spectrometry of peptides identified from male extracts (BRD-1 = 97.6% males; BRD-1::GFP::3xFLAG = 98.5% males). Ven diagram (generated by https://bioinfogp.cnb.csic.es/tools/venny/) of unique peptides from BRD-1::GFP::3xFLAG (purple) and BRD-1 peptides (green) with common peptides in grey. A small number of proteins with known interactions with BRC-1-BRD-1 (unique) as well as sperm proteins that are likely non-specific interactions (common) are highlighted. The complete data set of non-specific peptides and peptides unique to BRD-1::GFP::3xFLAG are available in Supplemental File 4.

### Data availability

Strains and reagents are available upon request. JEL1197 and JEL1214 will be deposited at CGC. The authors affirm that all data necessary for confirming the conclusions of this article are represented fully within the article and its tables and figures. Supplemental data are deposited at figshare.

## Results and Discussion

### Conditional degradation of DPY-27

The development of the auxin-inducible degradation system in *C. elegans* has facilitated analyses of many biological processes, including meiosis, spermatogenesis, mating, and aging (Kasimatis, Moerdyk-Schauwecker, and Phillips 2018; Ragle et al. 2020; Zhang et al. 2015; Cahoon and Libuda 2021). The system requires the introduction of a short amino acid sequence, auxin inducible degron (AID), onto a target protein, expression of the plant F-box protein TIR1, and the presence of the plant hormone auxin (Nishimura et al. 2009). TIR1 interacts with endogenous SKP1 and CUL1 proteins to form an SCF E3 ubiquitin ligase complex. In the presence of auxin, TIR1 recognizes and binds the AID sequence, leading to ubiquitination and subsequent degradation of the AID-tagged protein (Gray et al. 1999; Ruegger et al. 1998). We inserted an AID sequence together with a MYC tag at the C-terminus of DPY-27 (abbreviated as *dyp:27::AID*), which is a component of the DCC essential for embryonic development in hermaphrodites (Plenefisch, DeLong, and Meyer 1989), and generated *dpy-27::AID* strains that also expressed TIR1 under the control of the *sun-1* or *mex-5* promoter, which drive expression in germ cell and during early embryogenesis when dosage compensation is established (Ashley et al. 2021; Zhang et al. 2015). Additionally, these strains contain a mutation in *him-8*, which leads to the production of male self-progeny (Hodgkin, Horvitz, and Brenner 1979) (Figure 1A). The AID tag does not interfere with DPY-27 function, as in the absence of auxin there was no increase in male self-progeny (or decrease in hermaphrodite self-progeny) between AID tagged and non-tagged strains, indicating that DPY-27::AID is functional (*sun-1p::TIR1* or *mex-5p::TIR1; dpy-27::AID; him-8* vs. *him-8)* (Figure 1C). In the presence of 1mM auxin beginning at the L1 larval stage, the viable progeny of *sun-1p::TIR1; dpy-27::AID; him-8* hermaphrodites were almost exclusively males and the rare surviving hermaphrodites were dumpy, suggesting that dosage compensation was efficiently disrupted (Figure 1B, C). Treatment of L4 larvae also resulted in the production of almost all males (∼95%). In contrast, strains expressing the *mex-5* driven TIR1 did not result in as effective killing of hermaphrodite progenies (Figure 1C). Therefore, all subsequent analyses were performed using strains expressing the *sun-1* promoter driven TIR1.

### DPY-27 depletion does not affect male meiosis or spermiogenesis

A previous study had shown that DPY-28, another component of the DCC, plays a role in meiosis, but this appears to be independent of its function in DCC and is unrelated to DPY-27 (Tsai et al. 2008). To our knowledge, the effects on meiosis and spermatogenesis have not been examined in *dpy-27* mutant males. To determine whether meiosis was affected in males isolated following DPY-27 degradation, we examined the progression of meiotic recombination and sperm quality. We first monitored meiotic double strand break (DSB) repair by examining the assembly and disassembly of the recombinase RAD-51 (Rinaldo et al. 2002) in the spatiotemporal organization of the *C. elegans* male germ line using antibodies against RAD-51 (Checchi et al. 2014; Colaiacovo et al. 2003). RAD-51 loads onto resected DSBs destined for repair by homologous recombination beginning in leptotene/zygotene (transition zone) and is largely removed by mid-late pachytene. We had previously shown that *him-8* mutant males had wild-type levels of RAD-51 foci throughout the germ line (Jaramillo-Lambert and Engebrecht 2010). Comparison of wild type, *him-8*, and *sun-1p::TIR1; dpy-27::AID; him-8* without and with auxin treatment revealed no differences, suggesting that DPY-27 does not play a role in the assembly or disassembly of RAD-51 on meiotic DSBs (Figure 2A). To determine whether DSBs are accurately processed into crossovers, we monitored GFP::COSA-1, a cytological marker of crossover precursor sites (Yokoo et al. 2012). Wild-type males mostly exhibit five COSA-1 foci in pachytene nuclei, one on each of the five pairs of autosomes but not on the single *X* chromosome (Checchi et al. 2014). This pattern was unaltered by degradation of DPY-27 (WT=4.99±0.30, *him-8*=4.97±0.20, *sun-1p::TIR1; dpy-27::AID; him-8 -* auxin=5.00±0.13, +auxin=5.02±0.17; Figure 2B). Together, these results suggest that dagradation of DPY-27 has no effect on DSB repair and crossover designation in male meiosis.

Following crossover designation and resolution, spermatocytes undergo two rounds of divisions to produce haploid spermatids. Additionally, post-meiotic differentiation, spermiogenesis or sperm activation, is a pre-requisite for fertilization. To examine sperm morphology and activation, we released spermatids from male worms and examined their morphology under differential interference contrast microscopy in *sun-1p::TIR1; dpy-27::AID; him-8* in the absence or presence of auxin. We observed no difference in the morphology of the spermatids (Figure 2C). We also examined sperm activation by releasing spermatids into a solution containing Pronase E, which has previously been shown to induce differentiation into spermatozoa (Singaravelu et al. 2011). Under both conditions the activation of spermatids achieved a level greater than 80% (Figure 2C).

To monitor the quality of sperm produced following DPY-27 depletion, we mated the resulting males to determine if they produced euploid sperm competent for fertilization. We used the *fog-2(q71)* mutants for these experiments to eliminate hermaphrodite spermatogenesis, rendering *XX* animals self-sterile (Schedl and Kimble 1988), so that the contribution of the male parent to embryonic lethality could be assessed unambiguously. We observed no difference between wild type, *him-8* and *sun-1p::TIR1; dpy-27::AID; him-8* in the absence or presence of auxin (Figure 2D). Together, these results indicate that degradation of DPY-27 by the AID system does not affect meiosis or spermiogenesis.

### Molecular and biochemical studies using males collected following DPY-27 degradation

We used the *dyp-27::AID* system to monitor gene expression in the presence and absence of auxin. We first examined the expression of *vit-2*, one of 6 *vit* genes that encode vitellogenin (yolk proteins). *Vit* genes are expressed in the intestine of adult hermaphrodites, where the corresponding proteins are synthesized and then transported from the intestine into maturing oocytes (Goszczynski et al. 2016; Grant and Hirsh 1999; Kimble and Sharrock 1983). We collected adult worms from wild type (99.8% hermaphrodites/0.2% males) and *sun-1p::TIR1; dpy-27::AID; him-8* in the absence (60% hermaphrodites/40% males) and presence of auxin (0.5% hermaphrodites/99.5% males), extracted total RNA and analyzed *vit-2* expression using quantitative RT-PCR. We observed a small reduction in the expression of *vit-2* in *sun-1p::TIR1; dpy-27::AID; him-8* worms isolated from cultures grown in the absence of auxin compared to wild type (1.18-fold reduction), even though there was a 1.67-fold reduction in the number of hermaphrodites (Figure 3A). This smaller than expected reduction in expression may be a consequence of the smaller body size of males compared to hermaphrodites, and/or due to inefficient collection of the male worms when a significant proportion of the population is hermaphrodites. On the other hand, we observed a 300-fold reduction of *vit-2* expression in *sun-1p::TIR1; dpy-27::AID; him-8* worms isolated from cultures grown in the presence of auxin compared to wild type (Figure 3A). These results are consistent with strong enrichment of males in our cultures.

We next examined the expression of a male-specific transcription factor *snpc-1*.*3. snpc-1*.*3* drives male piRNA expression and is expressed in the male germ line (Choi et al. 2021). We observed a 4-fold enrichment of *snpc-1*.*3* expression in *sun-1p::TIR1; dpy-27::AID; him-8* worms isolated from cultures grown in the absence of auxin and a 60-fold enrichment of *snpc-1*.*3* in the presence of auxin compared to wild type. Thus, our enrichment procedure facilitates analyses of sex-specific gene expression profiles.

Gene expression studies can be performed on a relatively small number of worms; however, biochemical analyses require more material. To determine the utility of our male enrichment system in biochemical analyses, we isolated large numbers of males following DPY-27 degradation from worms expressing wild-type BRD-1 or BRD-1::GFP::3xFLAG, a component of the *C. elegans* BRCA1-BARD1 (BRC-1-BRD-1) complex. BRC-1-BRD-1 is expressed throughout the gonad and regulates several aspects of meiotic recombination in both oogenesis and spermatogenesis (Boulton et al. 2004; Janisiw et al. 2018; Li, Hariri, and Engebrecht 2020; Li et al. 2018; Polanowska et al. 2006).

We used magnetic anti-FLAG beads to pull down proteins associated with BRD-1::GFP::3xFLAG from male whole worm lysates (see Materials and Methods). Mass spectrometry analyses of two independent pull downs from both BRD-1 and BRD-1:: GFP::3xFLAG lysates identified enrichment of sperm proteins (Figure 3B). While these sperm proteins are likely not specific interactors of BRD-1, their identification is consistent with the strong enrichment of male worms following DPY-27 degradation. In the BRD-1::GFP::3xFLAG pull downs we specifically identified its binding partner BRC-1, H2A, a known substrate of the mammalian BRCA1-BARD1 E3 ligase (reviewed in (Witus, Stewart, and Klevit 2021) and LET-70/UBC-2, an E2 which had previously been shown to interact with the complex during DNA damage signaling in hermaphrodite worm extracts (Polanowska et al. 2006) (Figure 3B; Supplemental File 4). These results are consistent with a role for BRC-1-BRD-1 E3 ligase activity in male meiosis.

In conclusion, we describe a method to collect relatively pure populations of males that will facilitate future studies investigating how sex influences physiology. In combination with the large number of available mutants, and continued improvement on the auxin induced degradation system (Divekar et al. 2021; Negishi et al. 2021; Hills-Muckey et al. 2021), investigators can easily manipulate sex and determine the molecular processes that are differentially regulated.

## ACKNOWLEDGEMENTS

We thank the Caenorhabditis Genetic Center, which is funded by NIH Office of Research Infrastructure Programs (P40 OD010440) for providing strains. We are grateful to the Engebrecht lab for thoughtful discussions. We also thank Michelle Salemi and the UC Davis Proteomics Facility for mass spectrometry analyses. This work was supported by National Institutes of Health GM103860 and GM103860S1 to JE.

